# Capture probe, metabarcoding, or shotgun sequencing: which reflects local vegetation best?

**DOI:** 10.1101/2025.10.06.680654

**Authors:** Nichola A. Strandberg, Lucas Dane Elliott, Dilli Prasad Rijal, Dorothee Ehrich, Youri Lammers, Aloïs N. Revéret, Nigel G. Yoccoz, Iva Pitelkova, Antony G. Brown, Tyler J. Murchie, Kathleen Stoof-Leichsenring, Inger Greve Alsos

## Abstract

1. Metabarcoding is the most widely applied method for studying plant communities using environmental DNA, with shotgun sequencing and capture probes being alternative methods. Any method’s ability to detect and correctly identify plant taxa may vary with DNA preservation, DNA reference library and the size of the local flora, making it difficult to compare results from different environments.
2. Here we compare these methods using lake surface-sediments from Northern Fennoscandia with the PhyloNorway genome skim reference library (1500 taxa) that includes almost all species of the regional flora. We also undertook vegetation surveys from around the lakes to estimate the true positive detection rate, identify false positive detections, and provide optimal filtering cut-off thresholds for the three methods.
3. Optimal filter cutoffs were one PCR replicate and 3 reads for metabarcoding, and 3.2×10e-6 and 5.6×10e-6 of queried reads, respectively, for the shotgun and capture probe data. Applying these thresholds, the rate of false positives was too high for reliable identification at species level based on shotgun (49%) and capture probes (62%), whereas it was low for metabarcoding (5-12%). All methods were reliable at genus and family levels after applying the optimal filtering thresholds (<4% false positives). Our results show that metabarcoding on average detects 2.1 times as many true positive taxa as shotgun sequencing, and 6.4 times as many taxa as capture probes. Proportions of filtered metabarcoding and shotgun reads were significantly related to abundance categories from vegetation surveys, but this was not the case for capture probe data.
4. We expect the false positive rate of shotgun sequencing to decrease with increasing genome completeness in the reference libraries, and be advantageous for highly degraded DNA with fragments too short for metabarcoding. At present, metabarcoding provides the highest detectability and taxonomic resolution for correct identification of vascular plants.

## Introduction

Sedimentary ancient DNA (*seda*DNA) analysis is a fast-growing approach in palaeoecology and has several benefits compared to traditional approaches, since it can detect more taxa than macrofossil analysis and pollen analysis (Garcés-Pastor et al., 2023), providing a powerful tool for assessing biodiversity dynamics over long, e.g., millennial timescales (Capo et al., 2023). Currently, the most widely applied *seda*DNA method is metabarcoding (Alsos et al., 2024; Von Eggers et al., 2024), but alternatives exist in shotgun sequencing and capture probes (Murchie, Monteath, et al., 2021).

When metabarcoding, a single locus, e.g. a taxonomic diagnostic fragment of the plant chloroplast DNA, is amplified through PCR (polymerase chain reaction) using specific primer pairs (Garcés-Pastor et al., 2023). PCRs can be repeated multiple times for each sample extract, thus improving the chances of detecting rare or degraded ancient DNA sequences (Ficetola et al., 2015). However, the PCR step may skew the abundances of DNA fragments (Garcés-Pastor et al., 2023). For example, for the commonly used universal vascular plant *trn*L g/h primers (Taberlet et al., 2007) some plant families like Poaceae and Cyperaceae generally show lower abundances than expected (Anonymous, 2018), possibly due to GC content, fragment length and complexity (Nichols et al., 2018). Although taxonomic resolution is limited in some families, e.g. Salicaceae and Cyperaceae (Sønstebø et al., 2010), this is the method with overall highest taxonomic resolution (Revéret et al., 2023), with 40-50% of taxa identified to species level (Garcés-Pastor et al., 2025; Julián-Posada et al., 2025; Rijal et al., 2021).

Shotgun sequencing, also known as metagenomics, is the non-targeted sequencing of the DNA extracted and does not use primers thus overcoming issues with PCR bias. Shotgun sequencing can take advantage of whole genome references, to maximise information, and has the added advantage of retaining deamination damage at the ends of DNA sequences, allowing the distinction between ancient and modern DNA (Dabney et al., 2013). At present, a major drawback of shotgun sequencing is that it cannot reliably identify taxa at species level, due to incomplete reference databases, particularly for taxonomic groups lacking complete reference genomes such as plants (Revéret et al., 2023). This can result in more false positives (FPs) since the typically short ancient DNA sequences may be matched to incorrect taxa within these incomplete reference databases (Elliott et al., 2025; Wang et al., 2021), and many false negatives (FNs) since whole genome reference libraries are currently incomplete. In addition, shotgun sequencing typically generates large amounts of data with a small fraction allocated to eukaryotes, and larger proportion to prokaryotes which often remains unidentifiable (Wang et al., 2021). Together, these limitations, high data volume with low target yield, and higher costs currently restrict the effectiveness and efficiency of shotgun sequencing for *seda*DNA studies compared to more targeted approaches.

Capture enrichment (also known as hybridisation capture, capture enrichment and target capture) uses a bait-set to bind to targeted DNA sequences. The binding affinity to targets can be adjusted during bait design by adjusting bait tiling density, sequence identity and overlap through bait clustering, soft-masking, and taxonomic binning to improve specificity, and during wet lab processing by varying the hybridisation temperature, allowing closely related taxa, deaminated DNA fragments, and DNA fragments with individual variations to be retained (Capo et al., 2023). To date it is the least applied *seda*DNA method and has mainly been used to target a single genus (Meucci et al., 2021; Schulte et al., 2021). A number of probe sets exist that target species across many vascular plant families; the 353 probe set targeting nuclear Angiosperm genes (Johnson et al., 2019), the OZBaits_CP set, targeting 20 plastid gene regions (Foster et al., 2024) and the PaleoChip, targeting *mat*K, *rbc*L and *trn*L of the chloroplast (Murchie, Kuch, et al., 2021). To our knowledge, only the latter probe set has been applied to ancient samples of several sites (Kjær et al., 2022; Murchie, Monteath, et al., 2021), so it is difficult to assess the method’s efficiency. Also, as most studies do not have independent data available that could be used to distinguish true from false positive identifications, the optimal filtering criteria for reliable identification is largely unknown.

Studying *seda*DNA records from Northern Fennoscandia allows for a unique comparison of the methods as the regional flora is well known and relatively small, with a total of ∼600 species (Elven et al., 2022) and 250–550 species per 75 x 75 km grid (Grytnes et al., 1999). Almost all vascular plant species in the region are included in the genome skim reference library PhyloNorway, which on average covers 60% of the genome (Alsos et al., 2020; Wang et al., 2021). Here, we applied all three methods using established protocols to analyse lake surface sediment samples, and we validated these methods by comparing the results to field observations of plant biodiversity around the lakes. We first use the vegetation data to define filtering criteria that balance the loss of true positives versus retaining false positives. We then compare the three methods in taxonomic resolution and ability to detect local vegetation, taking into account ranked abundance.

## Methodology

### Selection of lakes

Lake catchments were selected covering the major vegetation types and climate gradients in northern Fennoscandia (Figure 1, Table S1). They represent the northernmost pine forest, widespread birch forest, as well as the arctic-alpine heath. The lakes have small inflows and are undisturbed by human activities except for reindeer grazing and a conifer plantation within one catchment (Sierravannet).

**Figure 1.**
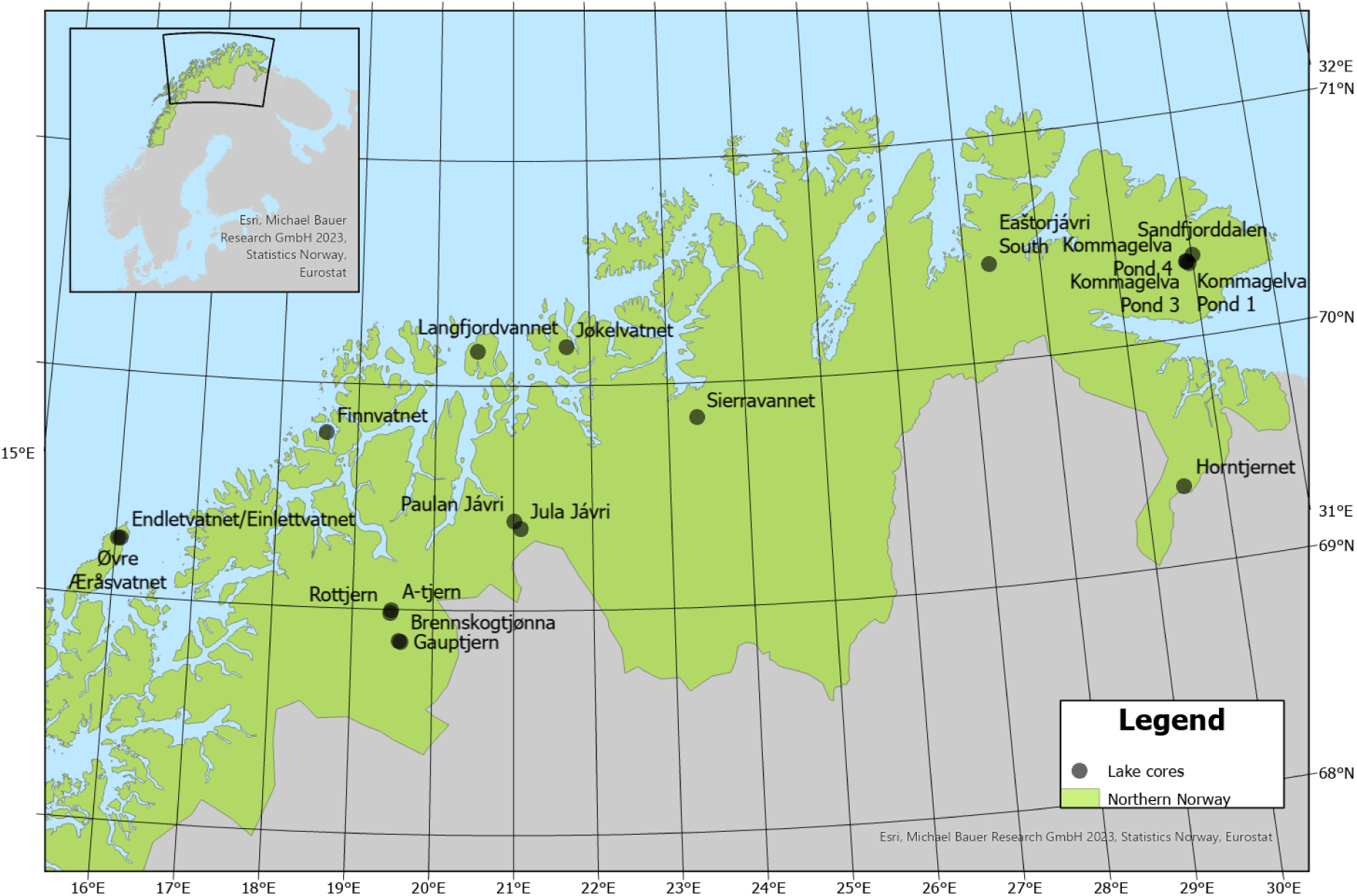
Location of lakes where vegetation surveys and DNA methods are compared.

**Figure 2.**
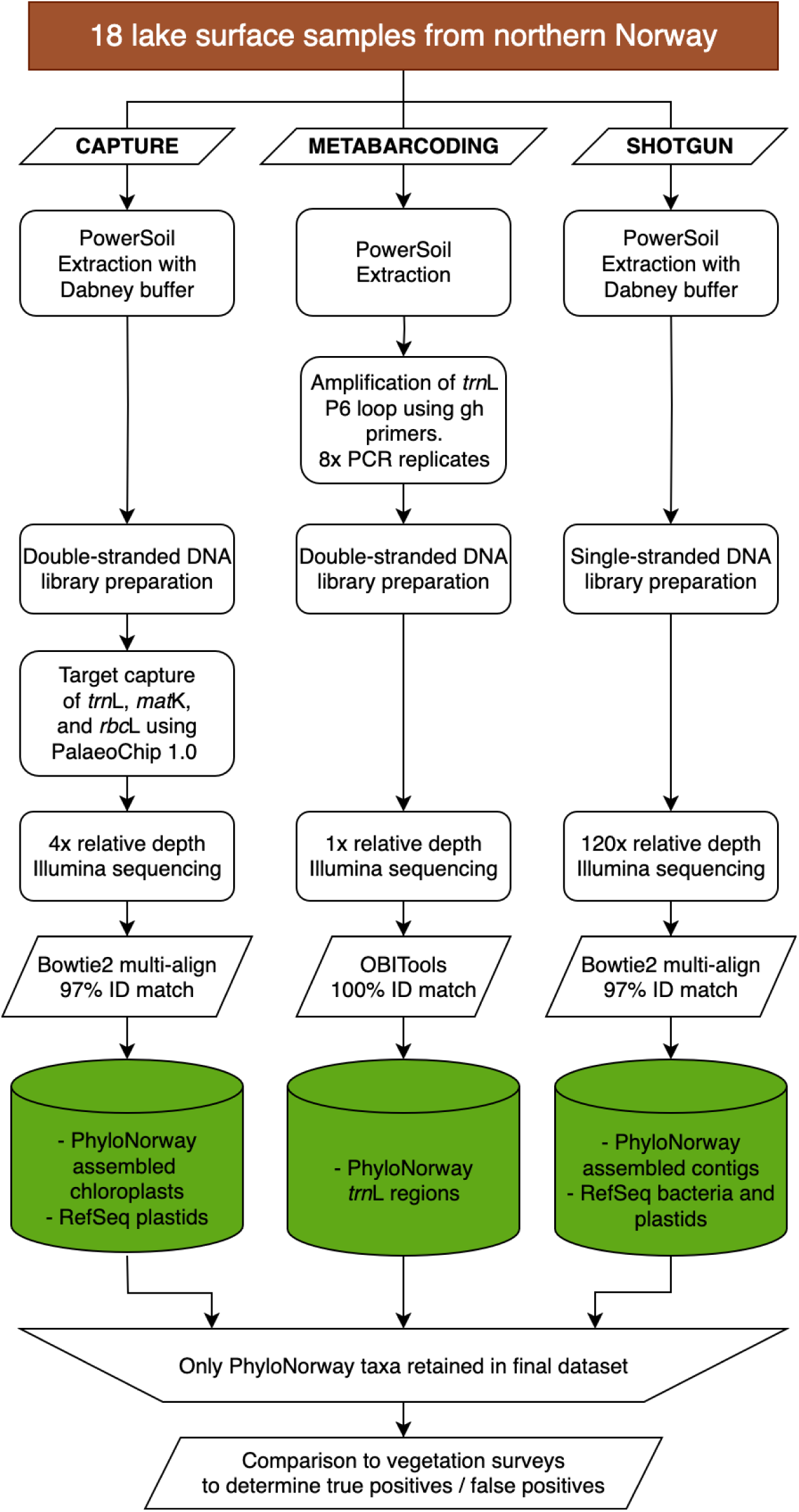
Workflow for the capture probe, metabarcoding and shotgun sequencing approaches.

### Vegetation surveys

Vegetation surveys recorded every plant taxon observed within 2m from the lakeshores. For the lakes included in Alsos et al. (2018), aquatic plants were surveyed from a boat using water binoculars and a long-handled rake. For the other lakes, the aquatic vegetation was surveyed by wading to collect and identify taxa. We noted plant taxa observed in the catchment >2 m away from the lakeshore, although this extended vegetation survey was not comprehensive. All plant vouchers are at the herbarium at The Arctic University Museum of Norway (TROM). Both terrestrial and aquatic plant taxa identified were given a ranked abundance score 1-4 with 1 being rare (only a few ramets), 2 being scarce (ramets occur throughout but at low abundance), 3 being common (common throughout but not the most abundant ones), and 4 being dominant (making up most of the biomass of the field, shrub or tree layer).

### Coring

Surface sediment were retrieved using either a Kajak 3 cm diameter mini gravity corer, a 5.9 cm diameter UWITEC USC 06000 corer, or a 4 cm diameter rod-operated Multisampler 12.42.01B, (Anonymous, 2018; Rijal et al., 2021). Eight samples were previously collected by (Anonymous, 2018), six samples by Rijal et al. (Rijal et al., 2021), one sample was used by both studies and three samples are new to this study. The cores were opened and subsampled in a dedicated ancient DNA laboratory at The Arctic University Museum of Norway in Tromsø.

### Capture probe

The DNA used for the capture probe analysis was extracted following a modified version of the Qiagen DNeasy PowerSoil PowerLyzer protocol as described by Rijal et al. (Rijal et al., 2021), which had an additional overnight centrifuge step to remove inhibitors (Supplementary Methods). An extraction control was processed alongside each group of 8 samples. The extracts were library prepared, along with a library control, following the double-stranded method (Meyer & Kircher, 2010) with dual-indexing modifications from (Kircher, 2012). The libraries were enriched using the PalaeoChip ArcticPlant-1.0 bait-set (Murchie, Kuch, et al., 2021). This bait-set primarily targets the plastid *trn*L (UAA) region from ∼2100 arctic vascular plant and bryophyte taxa, along with the full *rbc*L and *mat*K loci to increase the capture scope. The enriched libraries were then sequenced on an Illumina HiSeq 1500 at 2 x 90 bp at the Farncombe Metagenomics Facility (McMaster University, ON) (Supplementary Methods). The sample from lake Nesservatnet (NESS) only produced 3,763 read pairs and was excluded from further analysis.

### Metabarcoding

For metabarcoding, the DNA extraction, PCR, library cleaning and preparation, and sequencing protocol followed (Rijal et al., 2021). The 10 samples from (Anonymous, 2018) were re-extracted as these were initially only processed with six replicates. Nine extraction controls were included during the separate rounds of extractions. The *trn*L p6-loop region was amplified for the samples with the “g/h” primers (Taberlet et al., 2007), along with seven negative and seven positive PCR controls. Eight PCR replicates were generated for each sample. The library prepared samples were sequenced on an Illumina NextSeq 550 at 2 x 150 bp at the UiT Genomics Support Centre in Tromsø (Supplementary Methods).

### Shotgun sequencing

Shotgun extractions followed the protocol described for the capture probe analysis. As no DNA extracts remained, the same homogenized sediment samples were re-extracted (Supplementary Methods) with an extraction blank included for each group of eight samples. The samples were library prepared at the paleogenetic laboratories in the Alfred Wegener Institute (AWI) Helmholtz Centre for Polar and Marine Research in Potsdam, Germany, using a single-stranded library preparation method designed specifically for highly degraded ancient DNA (Gansauge et al., 2017; Gansauge & Meyer, 2013; Schulte et al., 2021). The library prepared samples, along with three extraction blanks, and one library blank, were sequenced on an Illumina NextSeq2000 at 2 x 100 bp at the sequencing facility at the Alfred Wegener Institute Helmholtz Centre for Polar and Marine Research, Bremerhaven, Germany (Supplementary Methods).

### Bioinformatics

#### Capture probe data

The sequenced reads were merged and adapter sequences removed using *fastp* v0.23.4 (Chen, 2023). Merged reads less than 30 bp were discarded as taxonomic resolution is poor for these short fragments (Pedersen et al., 2016). Reads were then mapped to a custom database using *bowtie2* with default parameters allowing for up to 1000 matches (Langdon, 2015). The custom database was constructed by compiling 1,541 assembled plastid genomes produced by the PhyloNorway project (Alsos et al., 2020) as well as the NCBI RefSeq plastid entries to allow for competitive mapping of potential cyanobacteria and algae sequences. Reads were assigned to the lowest common ancestor (LCA) using *ngsLCA* with a minimum edit distance proportion of 0.97 (Wang et al., 2022). Only those assignments to taxa represented in PhyloNorway with at least three reads were retained for further analysis.

#### Metabarcoding

The paired-end reads were merged, and adapter sequences were removed using SeqPrep (https://github.com/jstjohn/SeqPrep, v1.2). Reads were further processed using the OBITools pipeline (Boyer et al. 2016) largely following the protocol detailed in (Rijal et al., 2021). Taxonomic annotation was performed using a reference database of 1,541 P6-loop reference sequences from PhyloNorway (Alsos et al., 2020). Reads were assigned with 100% identity to a reference sequence and taxa were only retained with a minimum of three reads using a custom R script (https://github.com/Y-Lammers/MergeAndFilter).

#### Shotgun data

The sequenced reads were merged and adapter sequences removed using *fastp* v0.23.4 (Chen, 2023) with a minimum length of 30 bp. Taxonomic annotation was performed by *bowtie2* with default parameters allowing up to 1000 matches to a custom database (Langdon, 2015). This database was constructed by compiling the 1,541 partially assembled genome skims from PhyloNorway (Wang et al., 2021) as well as RefSeq bacteria and plastid entries to allow competitive mapping. Reads were assigned to the lowest common ancestor (LCA) using *ngsLCA* with a minimum edit distance proportion of 0.97 (Wang et al., 2022). Only those assignments to taxa represented in PhyloNorway with at least three reads were retained for further analysis.

### Comparison of DNA with vegetation surveys

The DNA detections that matched to the vegetation surveys at the site level were designated as true positives (TP). We then matched the remaining DNA detections with a list of plant taxa from the regional checklist for Northern Norway (Often & Alm, 1996) and designated these matches as regional flora (RF). These were not used further as we could not determine if they represented true or false positives. The detections with no match to either the vegetation surveys or regional flora were deemed false positives (FP). Designations were performed at the species, genus, and family taxonomic levels.

### Cutoff thresholds

Two approaches were used to set a minimum read threshold for a taxon to be retained in the molecular data. Both are based on the read proportion of a taxon relative to the total reads queried for that sample and aim to minimize the proportion of FPs relative to the total amount of TPs and FPs in the dataset. The first threshold is determined by finding the first local minimum value under 5% for FP/(FP+TP) while the second takes the absolute minimum value before the proportion reaches zero as all FP are discarded. These calculations were performed on the level of family, genus, and species, but thresholds were set using genus-level values.

### Read abundance and vegetation abundance

We performed an ordinal logistic regression using both the proportion of DNA reads and absolute read numbers (after applying optimal filtering) to predict the abundance categories for each taxon, and the transitions between the four abundance categories. This analysis was carried out at genus and species level. Furthermore, the relationship was examined using log, square root and double square root transformations. Models were fitted using the package *ordinal* (Christensen, 2012) in R version 4.4.3 with the clmm function (i.e. using a cumulative link model). Lake was included as random effect to account for differences in the vegetation survey ranked abundance estimations. For each sedaDNA method and transformation, we tested whether a flexible or equidistant model fitted the data better by comparing them with a likelihood ratio test. The fit of the models using different transformations of read proportions or numbers was compared with Akaike’s information criterion (AIC).

## Results

### Taxa detected in vegetation surveys

A total of 347 unique taxa were identified across all 18 lakes for both the 2 m and extended vegetation surveys. These taxa included 280 species belonging to 159 genera and 59 families (Table S2). The highest number of taxa were counted at Øvre Æråsvatnet (112), whereas only 33 species were counted at Kommagelva Pond 3.

### Sequencing results

From the capture probe samples, a total of 24,386,721 read pairs were obtained with an average of 1,274,382 ± 634,466 read pairs per sample (Table S3). The two extraction controls and library blank totalled 17,426 read pairs. After adapter trimming and merging, 19,705,695 (77.3%) and 9,216 (52.9%) sequences were retained for the samples and controls respectively. Four taxa were identified in the negative controls with >10 reads: Pinaceae/*Pinus* with 835 reads, Triticeae/*Triticum* with 26 reads, *Myrica* with 18 reads, and *Sparganium* with 15 reads. As Pinaceae/*Pinus* and *Myrica* do not appear in any samples from the extraction group (Table S4), we do not believe their presence in the dataset to be a result of contamination. *Triticum* appears in five samples from the extraction group and *Sparganium* appears in one sample, but as these are both designated regional flora (RF), their presence does not impact the evaluation of capture probe’s performance.

From the metabarcoding samples, a total of 7,923,720 read pairs were obtained with an average of 381,629 ± 88,837 per sample across eight PCR replicates. The nine extraction controls totalled 291,138 reads. After merging and processing through the OBITools pipeline, a total of 5,854,199 (76.7%) reads were retained and matched to a barcode. Across the nine extraction controls, 14 taxa were identified with six of them occurring in the samples from the same extraction group (Table S4). Four of these barcodes are only at family-level resolution and do not impact analyses, while the two others, *Cannabis* and *Pinus* are both designated regional flora at their respective sites and do not impact performance evaluation.

From the shotgun samples, a total of 904,790,751 read pairs were obtained with an average of 50,266,152 ± 13,462,696 read pairs per sample. The three extraction negative controls and two library blanks totalled 357,831 read pairs. After adapter trimming, merging, and filtering, a total of 699,592,884 (67.2%) and 15,675 (4.4%) sample and control sequences were retained respectively; only two taxa were identified in the negative controls with >10 read counts: *Avena* (oat) with 32 reads and *Triticum* (wheat) with 12 reads (Table S4). Both appear as only regional flora in the dataset and do not impact performance evaluation.

### Optimal read filtering threshold

At the species level, the number of false positives at relaxed filtering was high for both capture probe and shotgun sequencing (Table 1, Figure 3a), and it remained high (>30%) even with more stringent filtering criteria, making these methods unreliable at the species level (Table 1, Figure 3b). Thus, for capture probe and shotgun sequencing, the larger number of genus level identifications were used for more robust filtering estimates. The controls were not used for optimising the filtering threshold due to the relative lack of taxa detected. Metabarcoding detected 224 species at optimal filtering criteria, but the error rate was still relatively high (12%). This was partly caused by 17 fern detections that are known to appear in the gametophyte stage outside the range of the sporophyte (Brock, 2025). Disregarding ferns, the error rate for metabarcoding at species level was 5%.

**Table 1.** Table of false positive (FP) and true positive (TP) taxa detections and unique taxa retained at the genus- and species-level at different cutoff thresholds for each sequencing method. The cutoff thresholds are in order of increasing strictness starting with the initial base cutoff of three reads. The FP proportion is calculated by FP/(FP+TP) disregarding regional flora identifications.

| Method | Capture | Capture | Capture | Metabarcoding | Metabarcoding | Metabarcoding | Shotgun | Shotgun | Shotgun |
| --- | --- | --- | --- | --- | --- | --- | --- | --- | --- |
| Cutoff | 3 reads | 5.6x10e-6 | 1.5x10e-5 | 3 reads | 3.0x10e-5 | 3.4x10e-4 | 3 reads | 3.2x10e-6 | 2.18x10e-5 |
| FP genera detection proportion | 0.12 | 0.03 | 0.02 | 0.01 | 0.01 | 0.00 | 0.35 | 0.04 | 0.02 |
| FP genera detections | 11 | 2 | 1 | 5 | 4 | 0 | 371 | 7 | 1 |
| TP genera detections | 81 | 62 | 40 | 399 | 365 | 220 | 696 | 189 | 63 |
| FP genera | 10 | 2 | 1 | 1 | 1 | 0 | 108 | 5 | 1 |
| TP genera | 32 | 23 | 16 | 81 | 80 | 63 | 129 | 72 | 33 |
| FP species detection proportion | 0.74 | 0.62 | 0.33 | 0.12 | 0.10 | 0.07 | 0.79 | 0.49 | 0.37 |
| FP species detections | 49 | 23 | 4 | 30 | 23 | 10 | 1859 | 103 | 16 |
| TP species detections | 17 | 14 | 8 | 224 | 212 | 134 | 482 | 109 | 27 |
| FP species | 20 | 7 | 2 | 8 | 7 | 5 | 503 | 64 | 15 |
| TP species | 5 | 4 | 2 | 47 | 47 | 36 | 160 | 48 | 17 |

**Figure 3.**
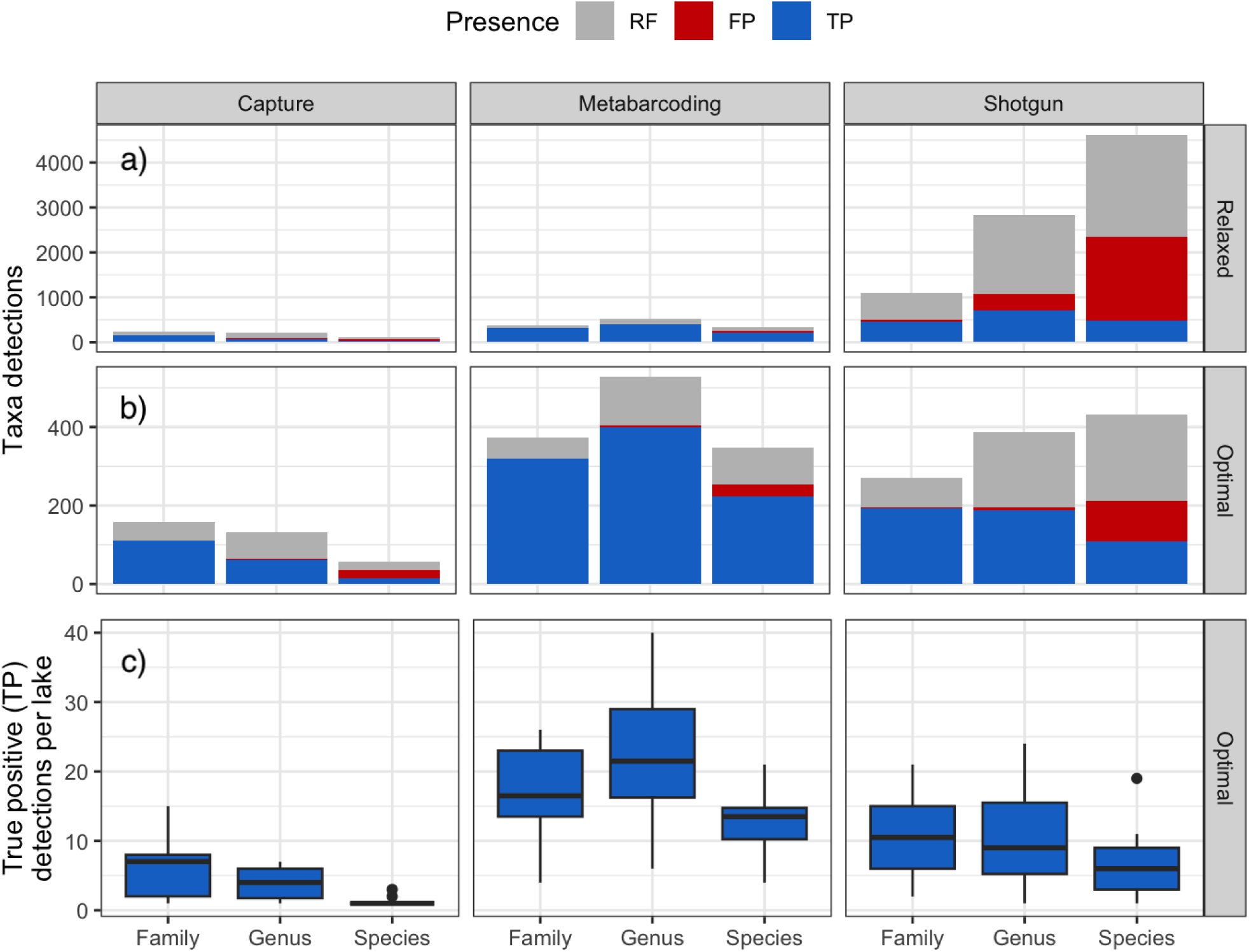
The count of true positive (TP), false positive (FP), and regional flora (RF) taxa identified by capture probe, metabarcoding, and shotgun methods. The values are displayed at the taxonomic levels of family, genus, and species for a) a relaxed cut-off of 3 reads for all methods; b) optimal cut-off of 3 reads for metabarcoding and 3.2×10e-6 and 5.6×10e-6 of reads for shotgun, and capture respectively, and c) number of true positive taxa detected at each lake after optimal filtering. A sequence identified at species- or genus-level is also included in higher taxonomic levels.

At the genus level, the total number of taxa detections (FP and TP) under the base filtering of 3 reads was highest for shotgun, but it had also the highest error rate (Table 1). Increasing the filtering stringency reduces both the total number of detections and the proportion of false positives across all methods, with a genus level error rate of 3% for capture probes, 1% for metabarcoding, and 4% for shotgun (Table 1). When applying optimal filtering level, the detection of TP genera was 2.1 times high for metabarcoding than shotgun, and 6.4 times higher for metabarcoding than capture probes (Table 1). These numbers do not take into account the 41 times deeper sequencing depth for shotgun than capture probes and 131 times deeper for shotgun than metabarcoding. When normalized by sequencing depth, capture probes detected 8 times more vascular plant genera than shotgun sequencing, and metabarcoding detected 26 times more than capture probes and 256 times more than shotgun sequencing. At the family level, the false positive rate was low for all three methods even at relaxed filtering (Figure 3a and b). The mean number of taxa detected at all three taxonomical levels was highest for metabarcoding, followed by shotgun sequencing, and lowest for capture probes (Figure 3c).

### Ability to detect local vegetation

Since shotgun and capture probe had high error rates at the species level even after our stringent filtering approach, we hereafter compare results at the genus level and only include true positives. Metabarcoding identified the greatest proportion of genera within our 2m vegetation surveys, and the detection rate was greater for the genera with higher ranked abundances (Figure 4). Rare taxa were detected at a rate of 2.5%, 7.8%, and 23% out of all possible observations by capture probe, shotgun, and metabarcoding methods respectively. In contrast, dominant genera were detected at a rate of 22%, 49%, and 77% respectively. The proportion of genera detected by all three methods was low, especially for less dominant taxa (Figure 4). Metabarcoding showed the greatest overlap with shotgun sequencing, while capture probe sequencing detected fewer genera overall. Further, there were more genera uniquely detected in metabarcoding (16–36%) than shotgun (3–5%) and capture probe (∼1–2%). At all ranked abundance categories, metabarcoding identified a greater proportion of genera than both metagenomic methods while shotgun outperformed capture probe.

**Figure 4.**
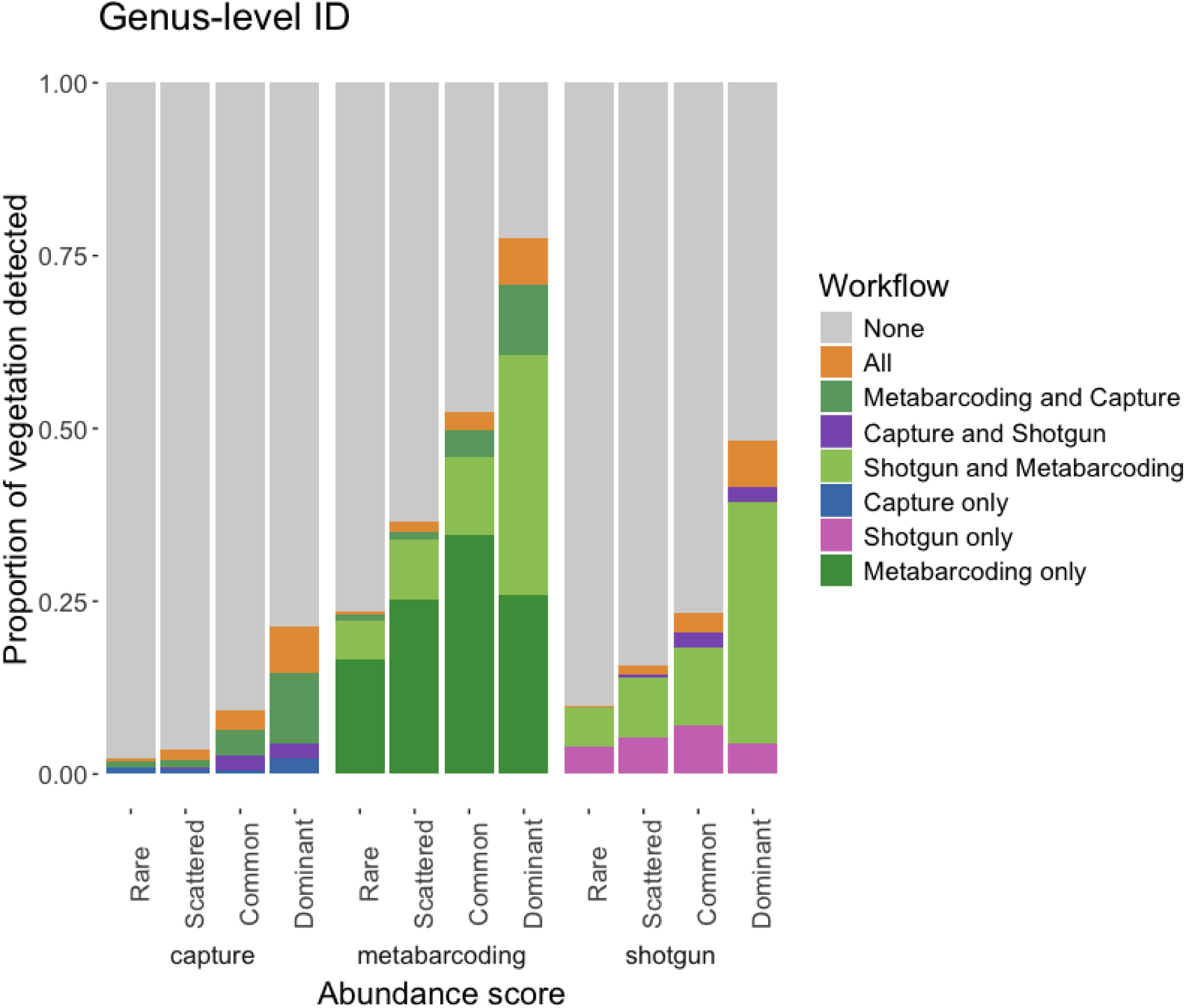
The proportion of 2 m vegetation survey taxa with a DNA match at the genus level grouped by DNA method detected by and ranked abundance. The DNA data shown are those done after the final optimal filtering.

### Correlation between sequence reads and the abundance in the vegetation

The proportion of reads for TPs at the genus level increased with abundance across taxa detected by metabarcoding (median proportion of reads: 0.00048, 0.00092, 0.00223, and 0.02105 for rare, scattered, common, and dominant genera, respectively). For shotgun sequencing there was a similar overall trend (median proportion of reads: 0.01034, 0.00690, 0.01153 and 0.02837 for rare, scattered, common, and dominant genera, respectively), whereas for capture probe data the relationship was less clear as the highest proportion of reads was recorded for scattered taxa (median proportion of reads:0.01233, 0.07030, 0.02580, 0.04468; Figure 5).

**Figure 5.**
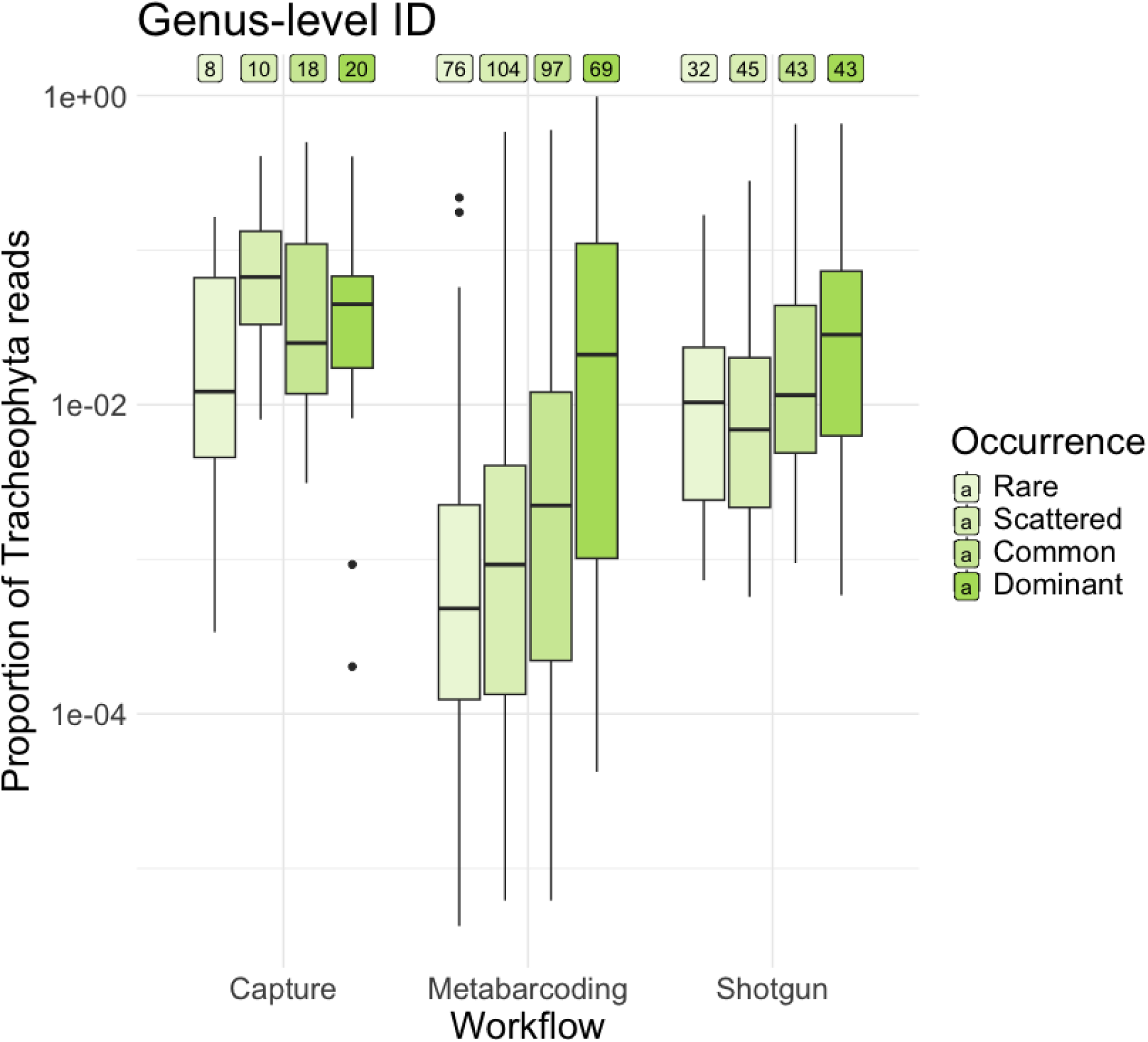
Proportions of Tracheophyta reads of the true positives at the genus level of the best estimated cutoff approach plotted according to vegetation survey ranked abundance. The horizontal line shows the median of the data and the boxes show the interquartile range. The numbers above the boxplots represent the number of genus observations within each category of plant abundance in vegetation surveys.

The ordinal regressions showed a significant relationship of the proportion of reads obtained from metabarcoding to the abundance category of plants observed within 2m from the lake shore (supplementary table x). Comparing different transformations of the read proportions with AIC suggested that double square root transformation resulted in the best model at both the genus and species level. At the species level, a log transformation was also supported (ΔAIC = 1.43). For shotgun data, relationships between the proportion of reads and the abundance categories were also significantly positive, but the results from AIC regarding the different transformations were less clear. All transformations seemed equally adequate at the species level, while at the genus level only the model without transformation appeared less supported (ΔAIC = 4.17). For the capture probe data, the proportion of reads was not related to the abundance categories at the genus level, while there were too few TPs at the species level to evaluate the relationship. Results were similar when calculated for the number of reads (Suppl. Table Sxx).

## Discussion

Our findings show that shotgun sequencing and capture probes require stringent filtering, whereas metabarcoding has a lower initial rate of false positives. All methods reliably identified taxa at genus level after optimal filtering whereas only metabarcoding produced reliable species-level identifications. Metabarcoding identified a larger proportion of genera at each abundance category compared to other methods. Additionally, DNA reads from metabarcoding and shotgun sequencing reliably predicted vegetation abundance, suggesting a link between read frequency and plant biomass. Although we aimed to optimise each laboratory protocol, this did result in different extraction and library preparation protocols used, which could introduce biases in the results. The sequencing depth for each method also varies with the shotgun sequencing having the highest sequencing depth followed by capture probe and then finally metabarcoding.

### Filtering

It is difficult to assess if our optimal filtering threshold of 5.6×10e-6 of the vascular plant reads for capture probes is more or less strict than used in previous studies, as they used a different approaches to filtering (Kjær et al., 2022; Murchie, Monteath, et al., 2021), but our data shows that cutoff level has a strong impact on detections and error rate for this type of data.

Our results suggest that a threshold of 3 or 10 reads is sufficient for metabarcoding. This is similar to a threshold of 5 (Stoof-Leichsenring et al., 2022) to 10 reads has been used in many studies (Garcés-Pastor et al., 2025; S. Liu et al., 2024; Rijal et al., 2021; Wang et al., 2021). Some studies apply additional criteria based on the repeatabilities across PCR replicates (Rijal et al., 2021) or clustering approaches (Y. Liu et al., 2025). While the absolute number of reads would vary with sample size and sequencing depth, metabarcoding seems to be reliable with a general low filtering threshold, and due to the earlier study on threshold level, the metabarcoding community in general apply adequate filtering threshold.

For shotgun sequencing, our estimated optimal cut-off level of 3.2×10e-6 of total reads sequenced assigned is less stringent than used by others. Wang et al. (Wang et al., 2021) applied an initial minimum filter of 1% of *Viridiplantae* reads and further removed the taxa with a combined read percentage lower than 5.39%, which is the median value they observed for Arctic genera. Courtin et al. (Courtin et al., 2022)) eliminated any sequences with an abundance of less than 0.2% of total *Viridiplantae* reads. Liu et al. (S. Liu et al., 2024) retained only those taxa detected in at least two samples with a combined read count of ≥5 therefore retaining 94% of their reads, which is less stringent than our filtering. Kjær et al. (Kjær et al., 2022) was more strict as they filtered out low-abundance taxa at the genus level by setting a threshold at half the median read count and retained taxa found in at least 3 samples. Thus, our use of vegetation data as an estimate of true and false positives may serve as a guideline for future studies.

For all three methods, the number of FPs is underestimated, as we only matched the sequences to PhyloNorway. While using global reference libraries as EMBL may increase detection, it may also inflate the FPs (Alsos et al., 2022). In cases where the local flora is less well covered in reference libraries or the reference library includes more erroneous annotated sequences, we advocate using a stricter cut-off level. As shotgun uses the whole genome, the effect of incomplete or erroneous annotated sequences will be considerably higher than for capture probes. Another source of potential FPs in the shotgun data is exogenous DNA within the reference databases. Even a well curated database like PhyloNorway (Alsos et al., 2020; Wang et al., 2021) contains endophytes such as bacteria and fungi (Elliott et al., 2025; Oskolkov et al., 2025), though increased reference material for these endophytes may reduce the false positives rate for shotgun data. Given the importance of filtering for biodiversity estimates, transparent reporting of these filtering criteria is crucial for reproducibility and meaningful comparisons.

### Methods performance

Our capture samples had on average 16 vascular plant taxa, thus lower than the average of 48.9 found by (Murchie, Monteath, et al., 2021) in their study from Pleistocene-Holocene transition permafrost samples and the 76.2 taxa per sample recorded by (Kjær et al., 2022). While capture probe is expected to give ∼1000× more ‘on-target’ DNA proportionally than comparable shotgun sequencing libraries (More et al., 2025), our capture identification was lower than the number of taxa found in shotgun analyses of the same samples. Thus, we suspect that the capture hybridization may not have worked optimally. One possible reason for the poor capture probe performance is that the Dabney extraction method, optimized for ultra-short DNA fragments (due to its high guanidine and isopropanol buffer), may be less effective for modern lake DNA. Additionally, degradation due to long transport time to Canada, issues such as enzyme/reaction failures during lab preparation or suboptimal bait hybridization could have degraded the performance. Further, we have noted that lake sediments may yield less DNA captured than permafrost samples using the PalaeoChip Arctic v1.0 Plant bait-set (Murchie pers. obs.). Further optimizations to the taxonomic breadth and genomic depth of an arctic plant bait set beyond *rbc*L, *mat*K, and *trn*L are recommended for future capture research if the focus is specifically on plants. Thus, overall, the method may have higher potential than found in our study.

The average number of TP and regional flora taxa found per sample using metabarcoding, 37.2, is higher than in the previous study of surface sediments from 11 lakes in the region (mean 19.7, (Anonymous, 2018), probably due to improved laboratory procedures. It is within the range of ancient samples in the region (20.6-65.5, n= 316 (Rijal et al., 2021) and surface samples in NE Siberia (15-54, n=32, (Niemeyer et al., 2017). It is considerably lower than 50-150 taxa per sample (n=705) found in the Alps, where a similar local DNA reference library is available (Garcés-Pastor et al., 2025), but that is probably due to the smaller species pool in the north. Thus, we conclude that the metabarcoding analyses here are representative for the method.

For shotgun analyses, the mean number of taxa per sample (25.5 when counting both TP and regional flora at family and genus level) was higher than the 8.13 and 17.5 taxa (at the genus level) detected by (Wang et al., 2021) and (Courtin et al., 2022) respectively. It was lower than (S. Liu et al., 2024) who found 40 taxa per sample and considerably lower than (Kjær et al., 2022) who found 76.21 taxa per sample, but had ten times higher sequencing depth. Given the high impact of both sequencing depth and cut-off level used on taxa richness reported, we conclude that our samples are within the range of expectation, and that they are representative for the method.

### Detectability and taxonomic resolution

Based on our results, metabarcoding appears to be currently the most effective method for detecting local vegetation from lake sediments. The addition of capture probe and shotgun sequencing provided only marginal improvements in taxon detection while also increasing false positive detections, and may not justify the increased laboratory time and costs for many studies. Similarly, a review of all studies that have detected aquatic macrophytes concluded that the detectability in general is higher using metabarcoding of the P6 loop than shotgun sequencing or capture probes (Revéret et al., 2023). Further, while shotgun and capture probes were not reliable at species level, metabarcoding was able to identify 47 species at <10% FP rate.

While the capture probe workflow annotated 3.4× more reads than shotgun sequencing relative to sequencing effort, the overall detection was still low. One limitation of the PalaeoChip bait set used in this study is that it targets the entire genes of *rbc*L, *trn*L, and *mat*K, which include highly conserved regions. These conserved regions are efficiently enriched but are often taxonomically uninformative, leading to plant identifications being assigned to higher taxonomic ranks such as phylum, class, or order (Murchie, Monteath, et al., 2021). However, a key advantage of targeting full genes is that it enables broader genomic coverage, which can support downstream analyses such as population genetics and the assessment of post-mortem DNA damage patterns. Another benefit of capture-based approaches is their compatibility with standard DNA barcoding markers, which in contrast to the non-standard minibarcode P6 loop, are available for the flora of most regions (https://boldsystems.org/).

While metabarcoding only explores a minor fraction of the plant genome and therefore has been argued to be less efficient than shotgun sequencing (Wang et al., 2021), our current comparison, as well as the majority of studies published earlier (Revéret et al., 2023), report considerable higher detection and taxonomic resolution for metabarcoding than shotgun data. Reliable species level identifications enables the reconstruction of both abiotic conditions such as temperature, pH, moisture, as well as biotic interactions as dispersal, pollination and mycorrhiza based on known species traits (Alsos et al., 2022; Garcés-Pastor et al., 2025). Also, metabarcoding is a more cost-efficient method as a much higher proportion of the sequences belong to target taxa, thus facilitating high time and space resolution analyses (Alsos et al., 2024). Furthermore, it is efficient in terms of bioinformatic requirements and relies on a reference library for only a small part of the genome.

Metabarcoding also has some down-sides compared to shotgun. If whole ecosystem reconstruction is the aim, additional analyses are required for other groups of organisms, by using mammal, fish, birds, or fungi specific primers (Capo et al., 2023), where the efficiency of each additional primer depends on the available reference data. Another downside is that metabarcoding relies on relatively long fragments (∼90 bases), making it less suitable for very degraded material. The oldest samples successfully analysed so far are Eemian lake sediments from arctic Canada (Crump et al., 2021), but for older samples or material from warmer regions, shotgun or capture probes may provide better results. Further, metabarcoding does not provide aDNA damage patterns due to the annealing of primers. Exploring damage patterns is desirable when focusing on species that are commonly found as contaminants in environmental and archaeological DNA studies (Smith et al., 2015; Weiß et al., 2015). It is also important when investigating caves or other non-water logged sediments where leaching is likely (Haile et al., 2007). However, no leaching/translocation has been documented for lake sediments (Sjögren et al., 2017), and they are generally regarded to provide reliable timelines for sedaDNA and many other determinants (Garcés-Pastor et al., 2023).

The low richness of shotgun may have resulted from both the high proportion of non-informative conserved genomic regions, and the incomplete reference libraries, preventing reliable identification at species level. When comparing sequencing depth of PhyloNorway to Kew plant genome sizes, PhyloNorway may have on average only approximately 60% genome coverage, which can introduce false positives by assigning reads to incorrect but closely related species. Increasing the amount of reference material should increase the true positive match for shotgun sequencing (Elliott et al., 2025), and thus projects that provide deep sequencing and/or fully assembled genomes for plants such as the Darwin Tree of Life (https://www.darwintreeoflife.org/) may considerably increase the potential for correct identifications. However, given the more than 300 000 species of vascular plants (Christenhusz & Byng, 2016), with relatively large and complex genomes, it will take time to complete. In regions with smaller floras, for example the Arctic islands, a complete reference database will be easier to obtain.

### Ability to record quantitative information

Our data showed that both metabarcoding and shotgun data show a clear increase in reads with increase in ranked abundance in the vegetation. For metabarcoding, similar results have also been found for a single lake compared to large sample vegetation surveys (Ataman et al., 2025) and for 200 x 1 m^2^ vegetation surveys compared to soil samples (Ariza et al., 2024). Thus, while PCR bias clearly does take place and for example grasses and sedges may be underrepresented (Anonymous, 2018), the overall quantitative pattern may be relatively robust. Similarly, although shotgun data may be biased in terms of unequal representation of the available reference genomes, the overall pattern remains robust. For both methods, one needs to take the taphonomical processes into account, which commonly causes over-representation of taxa growing along the streams and in the lakes. The lack of correlation of capture-probe data with plant abundance may have been due to the suboptimal performance of the method here.

## Conclusion

Each of the methods have their pros and cons; whilst capture probes and shotgun sequencing both have the advantage of displaying DNA damage patterns and thus may be the best choice if the focus is archaeological or non-water logged sediments, their ability to reflect plant diversity is limited by the lack of reliable information at species level. Metabarcoding is currently the most effective method for detecting vegetation given the current state of comparative reference databases, performing well at both the species and genus levels compared to shotgun sequencing and capture probe techniques. While shotgun and capture probe data can benefit significantly from filtering to reduce noise, metabarcoding tends to produce fewer false positives inherently. Additionally, both metabarcoding and shotgun methods show positive correlation between ranked taxonomic abundance and read counts, supporting their use in semi-quantitative analyses. Capture probes may increase the amount of target DNA sequenced, but further studies are needed to get a better understanding of its ability to capture quantitative patterns. While metabarcoding detected most true positives in our analysis, the high false positive rate associated with shotgun sequencing is expected to decline as more complete whole genome reference libraries become available. This advancement may make shotgun sequencing particularly advantageous for analysing highly degraded DNA, especially when fragment lengths are too short for successful metabarcoding.

## Acknowledgements

We thank Marie Føreid Merkel and Janine Klimke for laboratory assistance. The study was funded by Research Council of Norway grant 250963/F20 (to IGA, NGY, DE; supported DPR) for the ECOGEN project; The European Research Council (ERC) under the European Union’s Horizon 2020 research and innovation programme grant agreement No 819192 for the IceAGenT project (to IGA, AGB; supported YL); UiT and the ArcEcoGen Centre (supported LDE and NAS). Bioinformatic analyses were performed on resources provided by UNINETT Sigma2 - the National Infrastructure for High-Performance Computing and Data Storage in Norway.

## Data Availability

The raw capture probe, metabarcoding and shotgun sequencing data is deposited in the European Nucleotide Archive (ENA) under the accession code XXX. The identified taxa alongside the code for analyses and generating figures is available at https://github.com/salanova-elliott/Comparison_manuscript

## Author contributions

DPR and IGA designed the study. Vegetation surveys were conducted by NAS, LDE, DE, AGB, and IGA. Metabarcoding data was generated by DPR, ANR, YL, and IP. Shotgun data was generated by LDE and KS. Capture data was generated by IP and TM. Data was analysed by NAS, LDE, DPR, DE, NY, and IGA. The manuscript was written by NAS and LDE with feedback from all authors.

## References

Alsos, I. G., Boussange, V., Rijal, D. P., Beaulieu, M., Brown, A. G., Herzschuh, U., Svenning, J.-C., & Pellissier, L. (2024). Using ancient sedimentary DNA to forecast ecosystem trajectories under climate change. Philosophical Transactions of the Royal Society of London. Series B, Biological Sciences, 379(1902), 20230017.

Alsos, I. G., Lavergne, S., Merkel, M. K. F., Boleda, M., Lammers, Y., Alberti, A., Pouchon, C., Denoeud, F., Pitelkova, I., Pușcaș, M., Roquet, C., Hurdu, B.-I., Thuiller, W., Zimmermann, N. E., Hollingsworth, P. M., & Coissac, E. (2020). The treasure vault can be opened: Large-scale genome skimming works well using herbarium and silica gel dried material. Plants, 9(4), 432.

Alsos, I. G., Rijal, D. P., Ehrich, D., Karger, D. N., Yoccoz, N. G., Heintzman, P. D., Brown, A. G., Lammers, Y., Pellissier, L., Alm, T., Bråthen, K. A., Coissac, E., Merkel, M. K. F., Alberti, A., Denoeud, F., Bakke, J., & Null, N. (2022). Postglacial species arrival and diversity buildup of northern ecosystems took millennia. Science Advances, 8(39), eabo7434.

Anonymous. (2018). Plant DNA metabarcoding of lake sediments: How does it represent the contemporary vegetation. PloS One, 13(4), e0195403.

Ariza, M., Engelstad, M., Lieungh, E., Laux, M., Ready, J., Mauvisseau, Q., Halvorsen, R., & de Boer, H. J. (2024). Evaluating the feasibility of using plant-specific metabarcoding to assess forest types from soil eDNA. Applied Vegetation Science, 27(4). 10.1111/avsc.12806

Ataman, T. G., Lammers, Y., Alsos, I. G., Rijal, D. P., & Brown, A. G. (2025). Sedimentary DNA from lake depocenters maximizes detection of catchment vegetation. Communications Earth & Environment, 6(1), 1–10.

Brock, J. M. R. (2025). Effective dispersal of fern spore and the ecological relevance of zoochory. Biological Reviews of the Cambridge Philosophical Society, 100(5), 2116–2130.

Capo, E., Barouillet, C., & Smol, J. P. (Eds.). (2023). Tracking environmental change using lake sediments: Volume 6: Sedimentary DNA. Springer International Publishing.

Chen, S. (2023). Ultrafast one-pass FASTQ data preprocessing, quality control, and deduplication using fastp. iMeta, 2(2), e107.

Christenhusz, M. J. M., & Byng, J. W. (2016). The number of known plants species in the world and its annual increase. Phytotaxa, 261(3), 201.

Christensen, R. H. B. (2012). Ordinal: Regression Models for Ordinal Data. R Package Version 2011. 08-11.

Courtin, J., Perfumo, A., Andreev, A. A., Opel, T., Stoof-Leichsenring, K. R., Edwards, M. E., Murton, J. B., & Herzschuh, U. (2022). Pleistocene glacial and interglacial ecosystems inferred from ancient DNA analyses of permafrost sediments from Batagay megaslump, East Siberia. Environmental DNA (Hoboken, N.J.), 4(6), 1265–1283.

Crump, S. E., Fréchette, B., Power, M., Cutler, S., de Wet, G., Raynolds, M. K., Raberg, J. H., Briner, J. P., Thomas, E. K., Sepúlveda, J., Shapiro, B., Bunce, M., & Miller, G. H. (2021). Ancient plant DNA reveals High Arctic greening during the Last Interglacial. Proceedings of the National Academy of Sciences, 118(13), e2019069118.

Dabney, J., Meyer, M., & Pääbo, S. (2013). Ancient DNA damage. Cold Spring Harbor Perspectives in Biology, 5(7), a012567–a012567.

Elliott, L., Boyer, F., Lemane, T., PhyloAlps and PhyloNorway consortia, Alsos, I. G., & Coissac, E. (2025). Wholeskim: Utilising genome skims for taxonomically annotating ancient DNA metagenomes. Molecular Ecology Resources, e70001, e70001.

Elven, R., Bjorå, C. S., Fremstad, E., Hegre, H., & Solstad, H. (2022). Norsk Flora (Norwegian Flora). 8th edition. Samlaget.

Ficetola, G. F., Pansu, J., Bonin, A., Coissac, E., Giguet-Covex, C., De Barba, M., Gielly, L., Lopes, C. M., Boyer, F., Pompanon, F., Rayé, G., & Taberlet, P. (2015). Replication levels, false presences and the estimation of the presence/absence from eDNA metabarcoding data. Molecular Ecology Resources, 15(3), 543–556.

Foster, N. R., Jones, A. R., Serrano, O., Lafratta, A., Lavery, P. S., van Dijk, K.-J., Biffin, E., Gillanders, B. M., Young, J., Masque, P., Gadd, P. S., Jacobsen, G. E., Zawadzki, A., Greene, A., & Waycott, M. (2024). Environmental DNA identifies coastal plant community shift 1,000 years ago in Torrens Island, South Australia. Communications Earth & Environment, 5(1), 1–11.

Gansauge, M.-T., Gerber, T., Glocke, I., Korlević, P., Lippik, L., Nagel, S., Riehl, L. M., Schmidt, A., & Meyer, M. (2017). Single-stranded DNA library preparation from highly degraded DNA using T4 DNA ligase. Nucleic Acids Research, 45(10), e79–e79.

Gansauge, M.-T., & Meyer, M. (2013). Single-stranded DNA library preparation for the sequencing of ancient or damaged DNA. Nature Protocols, 8(4), 737–748.

Garcés-Pastor, S., Heintzman, P. D., Zetter, S., Lammers, Y., Yoccoz, N. G., Theurillat, J.-P., Schwörer, C., Tribsch, A., Walsh, K., Vannière, B., Wangensteen, O. S., Heiri, O., Coissac, E., Lavergne, S., van Vugt, L., Rey, F., Giguet-Covex, C., Ficetola, G. F., Karger, D. N., … Alsos, I. G. (2025). Wild and domesticated animal abundance is associated with greater late-Holocene alpine plant diversity. Nature Communications, 16(1), 3924.

Garcés-Pastor, S., Nota, K., Rijal, D. P., Liu, S., Jia, W., Leunda, M., Schwörer, C., Crump, S. E., Parducci, L., & Alsos, I. G. (2023). Terrestrial Plant DNA from Lake Sediments: Volume 6: Sedimentary DNA. In E. Capo, C. Barouillet, & J. P. Smol (Eds.), Tracking Environmental Change Using Lake Sediments (Vol. 21, pp. 275–298). Springer International Publishing.

Grytnes, J. A., Birks, H. J. B., & Peglar, S. M. (1999). Plant species richness in Fennoscandia: evaluating the relative importance of climate and history. Nordic Journal of Botany, 19(4), 489–503.

Haile, J., Holdaway, R., Oliver, K., Bunce, M., Gilbert, M. T., Nielsen, R., Munch, K., Ho, S. Y., Shapiro, B., & Willerslev, E. (2007). Ancient DNA chronology within sediment deposits: are paleobiological reconstructions possible and is DNA leaching a factor? Molecular Biology and Evolution, 24, 982–989.

Johnson, M. G., Pokorny, L., Dodsworth, S., Botigué, L. R., Cowan, R. S., Devault, A., Eiserhardt, W. L., Epitawalage, N., Forest, F., Kim, J. T., Leebens-Mack, J. H., Leitch, I. J., Maurin, O., Soltis, D. E., Soltis, P. S., Wong, G. K.-S., Baker, W. J., & Wickett, N. J. (2019). A universal probe set for targeted sequencing of 353 nuclear genes from any flowering plant designed using k-medoids clustering. Systematic Biology, 68(4), 594–606.

Julián-Posada, I., Gil-Romera, G., Garcés-Pastor, S., Heintzman, P. D., Gómez, D., Fillat, F., Moreno, A., Lara-Recuero, J., Bover, P., Montes, L., Sierra, A., Valero-Garcés, B., Alsos, I. G., & González-Sampériz, P. (2025). Neolithic pastoralism and plant community interactions at high altitudes of the Pyrenees, southern Europe. Communications Earth & Environment, 6(1), 1–10.

Kircher, M. (2012). Analysis of High-Throughput Ancient DNA Sequencing Data. In B. Shapiro & M. Hofreiter (Eds.), Ancient DNA: Methods and Protocols (pp. 197–228). Humana Press.

Kjær, K. H., Winther Pedersen, M., De Sanctis, B., De Cahsan, B., Korneliussen, T. S., Michelsen, C. S., Sand, K. K., Jelavić, S., Ruter, A. H., Schmidt, A. M. A., Kjeldsen, K. K., Tesakov, A. S., Snowball, I., Gosse, J. C., Alsos, I. G., Wang, Y., Dockter, C., Rasmussen, M., Jørgensen, M. E., … Willerslev, E. (2022). A 2-million-year-old ecosystem in Greenland uncovered by environmental DNA. Nature, 612(7939), 283–291.

Langdon, W. B. (2015). Performance of genetic programming optimised Bowtie2 on genome comparison and analytic testing (GCAT) benchmarks. BioData Mining, 8(1), 1.

Liu, S., Stoof-Leichsenring, K. R., Harms, L., Schulte, L., Mischke, S., Kruse, S., Zhang, C., & Herzschuh, U. (2024). Tibetan terrestrial and aquatic ecosystems collapsed with cryosphere loss inferred from sedimentary ancient metagenomics. Science Advances, 10(21), eadn8490.

Liu, Y., Lisovski, S., Courtin, J., Stoof-Leichsenring, K. R., & Herzschuh, U. (2025). Plant interactions associated with a directional shift in the richness range size relationship during the Glacial-Holocene transition in the Arctic. Nature Communications, 16(1), 1128.

Meucci, S., Schulte, L., Zimmermann, H. H., Stoof-Leichsenring, K. R., Epp, L., Bronken Eidesen, P., & Herzschuh, U. (2021). Holocene chloroplast genetic variation of shrubs (Alnus alnobetula, Betula nana, Salix sp.) at the siberian tundra-taiga ecotone inferred from modern chloroplast genome assembly and sedimentary ancient DNA analyses. Ecology and Evolution, 11(5), 2173–2193.

Meyer, M., & Kircher, M. (2010). Illumina sequencing library preparation for highly multiplexed target capture and sequencing. Cold Spring Harbor Protocols, 2010(6), db.prot5448.

More, K. D., Lebrasseur, O., Garrido, J. L., Seguin-Orlando, A., Discamps, E., Estrada, O., Tonasso-Calvière, L., Chauvey, L., Tressières, G., Schiavinato, S., Gibert, M., Padula, H., Chiavazza, H., Fernández, P. M., Guardia, N. M., Borges, C., Bertani, S., Contreras-Mancilla, J., Allccarima-Crisóstomo, D., … Orlando, L. (2025). Validating a target-enrichment design for capturing uniparental haplotypes in ancient domesticated animals. Molecular Ecology Resources, 25(7), e14112.

Murchie, T. J., Kuch, M., Duggan, A. T., Ledger, M. L., Roche, K., Klunk, J., Karpinski, E., Hackenberger, D., Sadoway, T., MacPhee, R., Froese, D., & Poinar, H. (2021). Optimizing extraction and targeted capture of ancient environmental DNA for reconstructing past environments using the PalaeoChip Arctic-1.0 bait-set. Quaternary Research, 99, 305–328.

Murchie, T. J., Monteath, A. J., Mahony, M. E., Long, G. S., Cocker, S., Sadoway, T., Karpinski, E., Zazula, G., MacPhee, R. D. E., Froese, D., & Poinar, H. N. (2021). Collapse of the mammoth-steppe in central Yukon as revealed by ancient environmental DNA. Nature Communications, 12(1), 7120.

Nichols, R. V., Vollmers, C., Newsom, L. A., Wang, Y., Heintzman, P. D., Leighton, M., Green, R. E., & Shapiro, B. (2018). Minimizing polymerase biases in metabarcoding. Molecular Ecology Resources. 10.1111/1755-0998.12895

Niemeyer, B., Epp, L. S., Stoof-Leichsenring, K. R., Pestryakova, L. A., & Herzschuh, U. (2017). A comparison of sedimentary DNA and pollen from lake sediments in recording vegetation composition at the Siberian treeline. Molecular Ecology Resources, 17(6), e46–e62.

Often, A., & Alm, T. (1996). Kryssliste for Nord-Norge (Checklist for Northern Norway). Polarflokken, 20(2), 165–168.

Oskolkov, N., Jin, C., Clinton, S. L., Guinet, B., Wijnands, F., Johnson, E., Kutschera, V. E., Kinsella, C. M., Heintzman, P. D., & van der Valk, T. (2025). Disinfecting eukaryotic reference genomes to improve taxonomic inference from ancient environmental metagenomic data. In bioRxiv (p. 2025.03.19.644176). 10.1101/2025.03.19.644176

Pedersen, M. W., Ruter, A., Schweger, C., Friebe, H., Staff, Richard A., Kjeldsen, K. K., Mendoza, M. L. Z., Beaudoin, A. B., Zutter, C., Larsen, N. K., Potter, B. A., Nielsen, R., Rainville, R. A., Orlando, L., Meltzer, D. J., Kjær, K. H., & Willerslev, E. (2016). Postglacial viability and colonization in North America’s ice-free corridor. Nature, 537, 45–49.

Revéret, A., Rijal, D. P., Heintzman, P. D., Brown, A. G., Stoof-Leichsenring, K. R., & Alsos, I. G. (2023). Environmental DNA of aquatic macrophytes: The potential for reconstructing past and present vegetation and environments. Freshwater Biology. 10.1111/fwb.14158

Rijal, D. P., Heintzman, P. D., Lammers, Y., Yoccoz, N. G., Lorberau, K. E., Pitelkova, I., Goslar, T., Murguzur, F. J. A., Salonen, J. S., Helmens, K. F., Bakke, J., Edwards, M. E., Alm, T., Bråthen, K. A., Brown, A. G., & Alsos, I. G. (2021). Sedimentary ancient DNA shows terrestrial plant richness continuously increased over the Holocene in northern Fennoscandia. Science Advances, 7(31), eabf9557.

Schulte, L., Bernhardt, N., Stoof-Leichsenring, K., Zimmermann, H. H., Pestryakova, L. A., Epp, L. S., & Herzschuh, U. (2021). Hybridization capture of larch (Larix Mill.) chloroplast genomes from sedimentary ancient DNA reveals past changes of Siberian forest. Molecular Ecology Resources, 21(3), 801–815.

Sjögren, P., Edwards, M. E., Gielly, L., Langdon, C. T., Croudace, I. W., Merkel, M. K. F., Fonville, T., & Alsos, I. G. (2017). Lake sedimentary DNA accurately records 20th Century introductions of exotic conifers in Scotland. The New Phytologist, 213(2), 929–941.

Smith, O., Momber, G., Bates, R., Garwood, P., Fitch, S., Pallen, M., Gaffney, V., & Allaby, R. G. (2015). Sedimentary DNA from a submerged site reveals wheat in the British Isles 8000 years ago. Science, 347(6225), 998–1001.

Sønstebø, J. H., Gielly, L., Brysting, A. K., Elven, R., Edwards, M., Haile, J., Willerslev, E., Coissac, E., Rioux, D., Sannier, J., Taberlet, P., & Brochmann, C. (2010). Using next-generation sequencing for molecular reconstruction of past Arctic vegetation and climate. Molecular Ecology Resources, 10(6), 1009–1018.

Stoof-Leichsenring, K. R., Huang, S., Liu, S., Jia, W., Li, K., Liu, X., Pestryakova, L. A., & Herzschuh, U. (2022). Sedimentary DNA identifies modern and past macrophyte diversity and its environmental drivers in high-latitude and high-elevation lakes in Siberia and China. Limnology and Oceanography, 67(5), 1126–1141.

Taberlet, P., Coissac, E., Pompanon, F., Gielly, L., Miquel, C., Valentini, A., Vermat, T., Corthier, G., Brochmann, C., & Willerslev, E. (2007). Power and limitations of the chloroplast trnL (UAA) intron for plant DNA barcoding. Nucleic Acids Research, 35(3), e14.

Von Eggers, J., Monchamp, M.-E., Capo, E., Giguet-Covex, C., Spanbauer, T., & Heintzman, P. (2024). Inventory of ancient environmental DNA from sedimentary archives: locations, methods, and target taxa V2. https://zenodo.org/records/13761348

Wang, Y., Korneliussen, T. S., Holman, L. E., Manica, A., & Pedersen, M. W. (2022). *ngs*LCA—A toolkit for fast and flexible lowest common ancestor inference and taxonomic profiling of metagenomic data. Methods in Ecology and Evolution, 13(12), 2699–2708.

Wang, Y., Pedersen, M. W., Alsos, I. G., De Sanctis, B., Racimo, F., Prohaska, A., Coissac, E., Owens, H. L., Merkel, M. K. F., Fernandez-Guerra, A., Rouillard, A., Lammers, Y., Alberti, A., Denoeud, F., Money, D., Ruter, A. H., McColl, H., Larsen, N. K., Cherezova, A. A., … Willerslev, E. (2021). Late Quaternary dynamics of Arctic biota from ancient environmental genomics. Nature, 600, 86–92.

Weiß, C. L., Dannemann, M., Prüfer, K., & Burbano, H. A. (2015). Contesting the presence of wheat in the British Isles 8,000 years ago by assessing ancient DNA authenticity from low-coverage data. eLife, 4. 10.7554/eLife.10005

