## Supplementary Information for "Capture probe, metabarcoding, or shotgun sequencing: which reflects local vegetation best?"

### Supplementary Methods

#### Capture probe extraction, capture and sequencing

The DNA used for the capture probe analysis was extracted following a modified version of the Qiagen DNeasy PowerSoil PowerLyzer protocol as described by Rijal et al. 2021. An additional centrifuge step where the PowerBead supernatant was added to 17.5 ml of Dabney binding buffer and spun at 4200 rpm for 20 hours overnight to remove inhibitors. An extraction control was processed alongside each group of 8 samples.

DNA extracts of 15 ng from the 18 lakes were enriched and sequenced at McMaster Ancient DNA Centre, Hamilton, Canada in 2019. The sequencing library was prepared, alongside one library control blank, using a double-stranded method (Meyer & Kircher, 2010) with modifications from (Kircher, 2012), and a modified end-repair reaction to account for the lack of uracil excision. Samples were purified after blunt-end repair with a QIAquick PCR Purification Kit (QIAGEN) (to maximally retain small fragments) and after adapter ligation with a MinElute PCR Purification Kit (QIAGEN). Libraries were quantified using the short amplification primer set with PCR duplicates to assess library preparation success and thereafter were uniquely dual indexed.
Libraries with a higher and lower amplification RFU values (relative to other libraries in the same indexing set) were removed after 11 and 14 PCR cycles respectively. All were

re-added for the final round of extension. The adapter-ligated, dual-indexed double-stranded DNA libraries were then enriched using the PalaeoChip ArcticPlant-1.0 bait-set, which had been designed in collaboration with Arbor Biosciences (Murchie, Kuch, et al., 2021). This bait-set targets ~2100 arctic vascular plant and bryophyte taxa, based on the databases available from Sønstebø et al. (Sønstebø et al., 2010), Soininen et al. (Soininen et al., 2015), and Willerslev et al. (Willerslev et al., 2014). The chloroplast locus *trn*L (UAA) is the primary target of these reference databases; additional full *trn*L loci from GenBank were added to the bait-set to augment some of the particularly short sequences (<50 bp) available in the original references. The full loci *rbc*L and *mat*K (each ~1500 bp) were also added where available to further increase the chloroplast targeting scope.

Hybridization and bait mixes were prepared to a desired concentration of 100 ng of baits per reaction. An indexed library input of 5 μL was combined with blOligos (blocking oligos which prevent the hybridization between library adapter sequences). The hybridization and bait mixes were pre-warmed to 60°C before being combined with the library- blOligo mixture. The final reaction was incubated for 45 hours at 60°C for bait-library hybridization. After the two-day hybridization, beads were dispensed (20 μL per reaction), washed three times with an equivalent volume of binding buffer per library, then resuspended in binding buffer and aliquoted into PCR strips. Baits were captured using 20 μL of the bead binding buffer suspension per library, incubated at 60°C for 2.5 minutes, finger vortexed and spun down, then incubated for another 2.5 minutes. Beads were pelleted and the supernatant (the non-captured library fraction) was removed and stored at −20°C as per (Klunk et al., 2019). The beads were resuspended in 180 μL of 60°C Wash Buffer X per tube and washed four times following the myBaits V4 protocol. Beads were eluted in 15 μL EBT, PCR reamplified for 12 cycles, then purified with MinElute columns following manufacturer’s protocols, and finally eluted in 15 μL EBT.

Total enriched DNA was quantified using the long amplification primer set with PCR duplicates. These values were used to calculate the dilution ratio for equimolar pooling, aiming for ~1,000,000 sequenced reads per library, but while maintaining a pool molarity of ≳200 pM for post-pooling purification as well as size-selection procedure. The pools were size-selected with gel excision following electrophoresis for molecules ranging between 150–600 bp. Gel plugs were purified using the QIAquick Gel Extraction Kit (QIAGEN), according to manufacturer's protocol, then sequenced on an Illumina HiSeq 1500 at 2 x 90 bp paired-end protocol at the Farncombe Metagenomics Facility (McMaster University, ON). The sample from lake Nesservatnet (NESS) only produced 3,763 read pairs and was excluded from further analysis.

#### Metabarcoding extraction, amplification and sequencing

For metabarcoding, the DNA extraction protocol followed that described by (Rijal et al., 2021). The extractions were carried out in a dedicated ancient DNA laboratory at The Arctic University Museum of Norway in Tromsø. The 10 samples from Alsos et al. (Alsos et al., 2018) were re-extracted as the extracts were depleted and the original metabarcoding procedure was carried out with only six replicates. In total nine extraction controls were included during the separate rounds of extractions.

The extracts were amplified with the "g/h" primers (Taberlet et al. 2007) that targets the vascular plant *trn*L p6-loop region located on the chloroplast. The primers were uniquely dual tagged with 8-9 bp tags, which allowed for pooling of multiple samples within the same sequencing library. The PCR reactions were carried out in a 40 µL final volume containing 4 µL of DNA extract, 1x Gold buffer, 1.6 U AmpliTaqGold Polymerase (Life Technologies), 2.5 mM $MgCl_2$, 0.2 mM dNTPs (VWR), 0.2 µM of each primer and 160 ng/µL Bovine Serum Albumin (Fisher). The PCR reaction started with an initial enzyme activation at 95 °C for

10 min, followed by 45 cycles of denaturation at 95 °C for 30 sec, annealing at 50 °C for 30 sec, and elongation at 72 °C for 1 min, with a final elongation step at 72 °C for 7 min. A total of eight PCR replicates were generated for each sample and control. In addition, nine and nine negative and positive control were included for the amplification.

After the amplification, the resulting PCR product was cleaned and pooled following the protocol described by Voldstad et al. (Voldstad et al. 2020). The resulting pools were converted into DNA libraries using the TruSeq PCR-free library kit with dual unique indexing. Library quantification was carried out via qPCR using the Illumina Library Quantification Kit (KAPA Biosystems) and a Prism 7500 Real-Time PCR System (Life Technologies). Each library was sequenced on ~10% of a flow cell on the Illumina NextSeq-550 platform (2 × 150 bp, mid-output mode) at the UiT Genomics Support Centre in Tromsø.

#### Shotgun extraction and sequencing

The same extraction protocol described for the capture probe analysis was used for the shotgun sequencing. As there were no DNA extracts remaining, another round of extraction on the same homogenized sediment subsamples was performed and subsequently processed for shotgun sequencing. Extraction blanks were included for each group of eight samples that were extracted.

Library preparation for shotgun sequencing was performed in the paleogenetic laboratories at the Alfred Wegener Institute (AWI) Helmholtz Centre for Polar and Marine Research in Potsdam, Germany, using a single-stranded library preparation method designed specifically for highly degraded ancient DNA (Gansauge et al., 2017; Gansauge & Meyer, 2013; Schulte et al., 2021). From the previously described extraction process, 15 ng DNA was used as a DNA input. In total three library batches containing 24 samples and three library blank controls were included.

Libraries were quantified using qPCR (Gansauge & Meyer, 2013): The qPCR setup contained 1x Maxima™ SYBR™ Green (Thermo Scientific, Germany), 0.2 µM IS7 and IS8, and 1 µL of the libraries diluted 1:20 with TET buffer in a final volume of 25 µL. The qPCR was run in Quant Studio 3 (Thermo Fisher Scientific) with the following settings: 95 °C for 10 min, followed by 40 cycles of 30 s at 95 °C, 30 s at 60 °C and 30 s at 72 °C. The fluorescence was acquired after each cycle and the amplification curve was used to estimate the needed number of amplification cycles during indexing PCR. Indexing PCR was performed in 10–14 cycles (for samples and blanks) depending on the library concentration with indexed P5 (5′−3′: AATGATACGGCGACCACCGAGATCTACACNNNNNNNACACTCTTTCCCTACACGACGCTCTT; IDT, Germany) and P7 (5′−3′: CAAGCAGAAGACGGCATACGAGATNNNNNNNGTGACTGGAGTTCAGACGTGT, IDT, Germany) primers using AccuPrime Pfx polymerase (Life Technologies, Germany) and 24 µL of each library. Amplificates were purified with the MinElute PCR Purification Kit (Qiagen, Germany) and library size distribution was checked on the Agilent TapeStation 4200 using the D1000 ScreenTape (Agilent Technologies, USA). The samples were pooled equimolarly, but with the negative control in a 10:1 ratio to achieve a final molarity of 20 mM. The 21 samples, three extraction blanks, and one library blank, were sequenced in paired-end mode (2 × 100 bp) on an Illumina NextSeq2000 device at the sequencing facility at The Alfred Wegener Institute Helmholtz Centre for Polar and Marine Research, Bremerhaven, Germany. One lake, Nesservatnet, failed to produce any sequences and was excluded from further analysis.

### Supplementary Figures

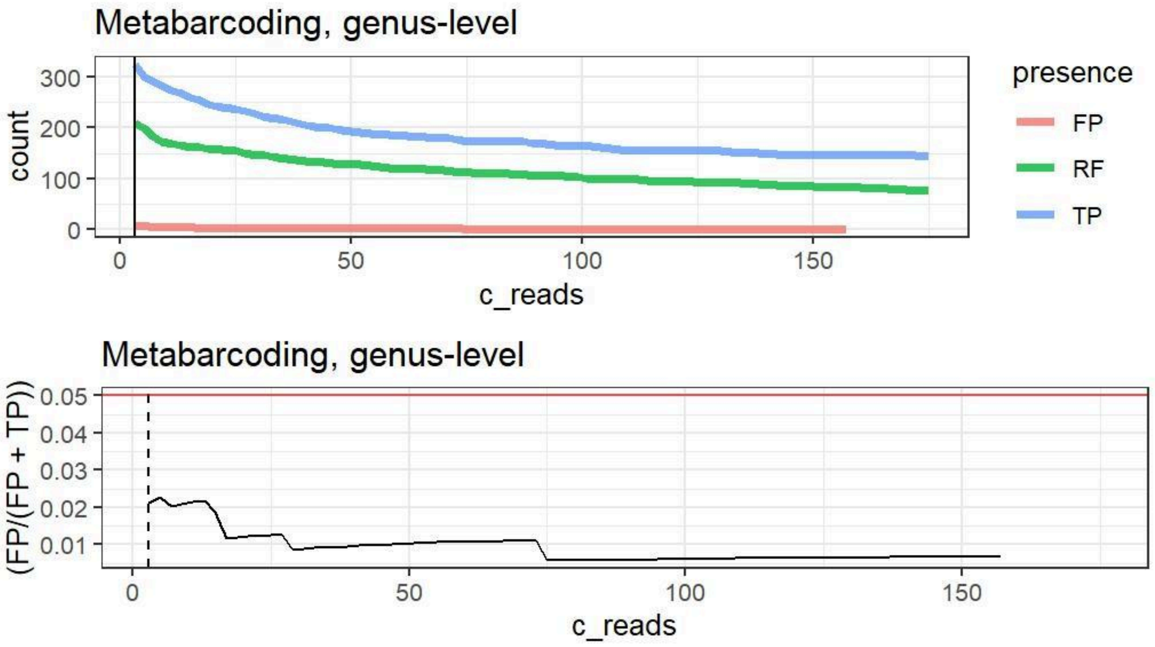


Figure S1. Top figure shows the number of true positive (TP), false positive (FP), and regional flora (RF) genera retained in the metabarcoding dataset at increasing read cutoffs. The bottom figure shows the proportion of FP genera in the dataset at increasing read cutoffs. The horizontal red line marks 0.05 proportion and the dashed vertical line marks the optimal cutoff threshold.

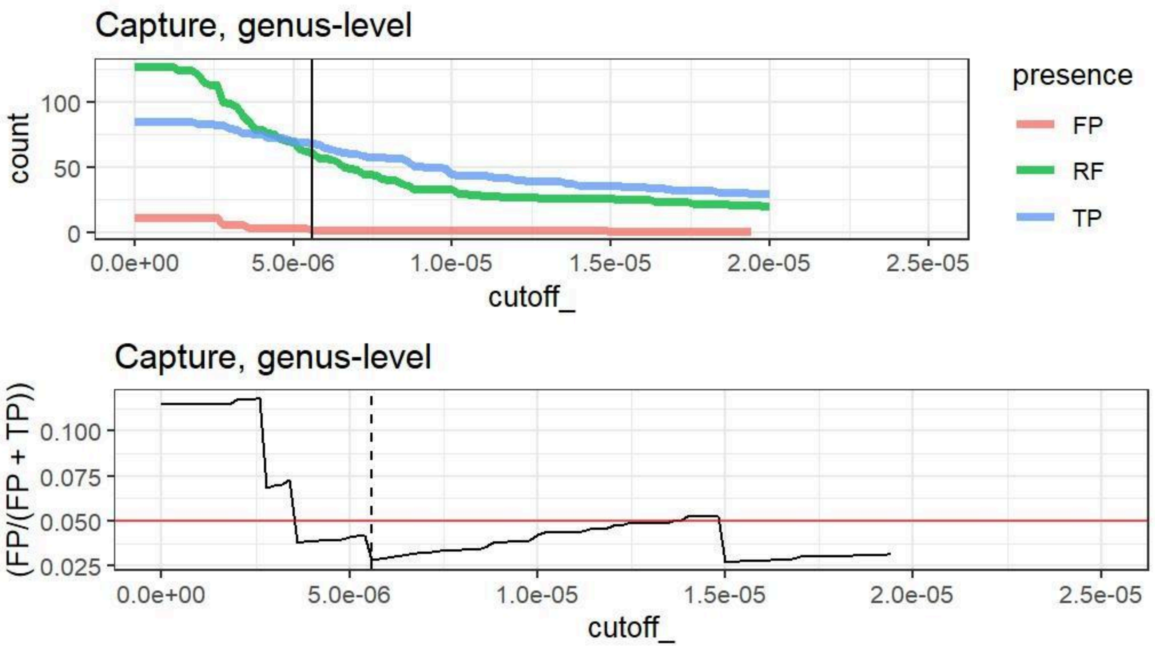


Figure S2. Top figure shows the number of true positive (TP), false positive (FP), and regional flora (RF) genera retained in the capture probe dataset at increasing read cutoffs. The bottom figure shows the proportion of FP genera in the dataset at increasing read cutoffs. The horizontal red line marks 0.05 proportion and the dashed vertical line marks the optimal cutoff threshold.

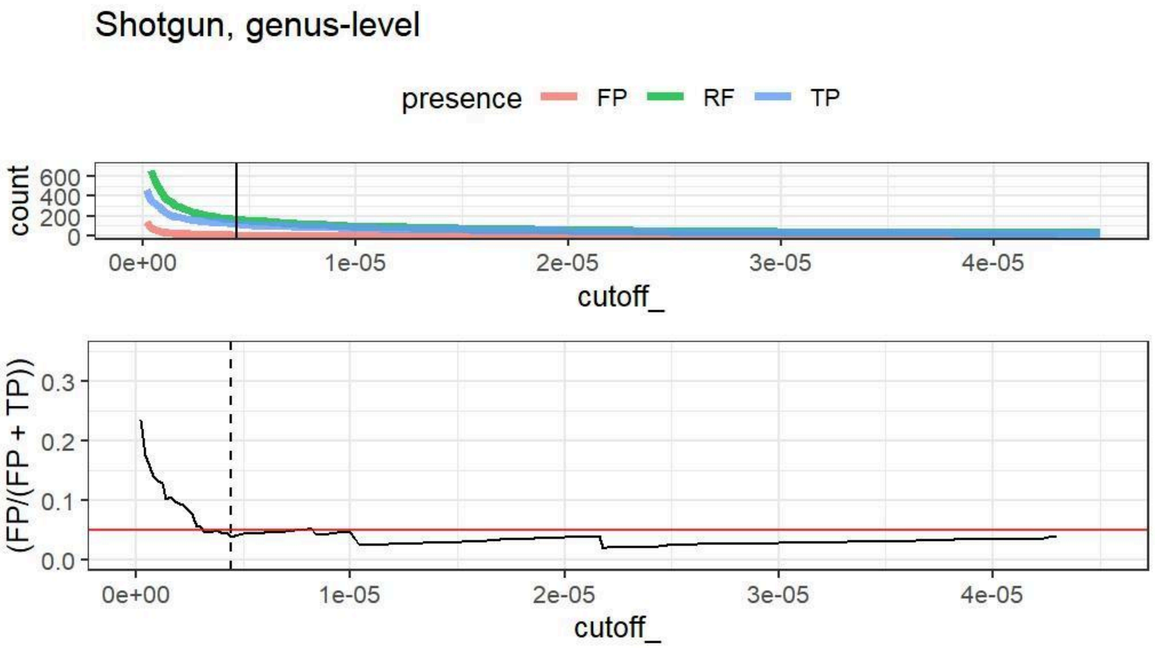

Figure S3. The top figure shows the number of true positive (TP), false positive (FP), and regional flora (RF) genera retained in the shotgun dataset at increasing read cutoffs. The bottom figure shows the proportion of FP genera in the dataset at increasing read cutoffs. The horizontal red line marks 0.05 proportion and the dashed vertical line marks the optimal cutoff threshold.

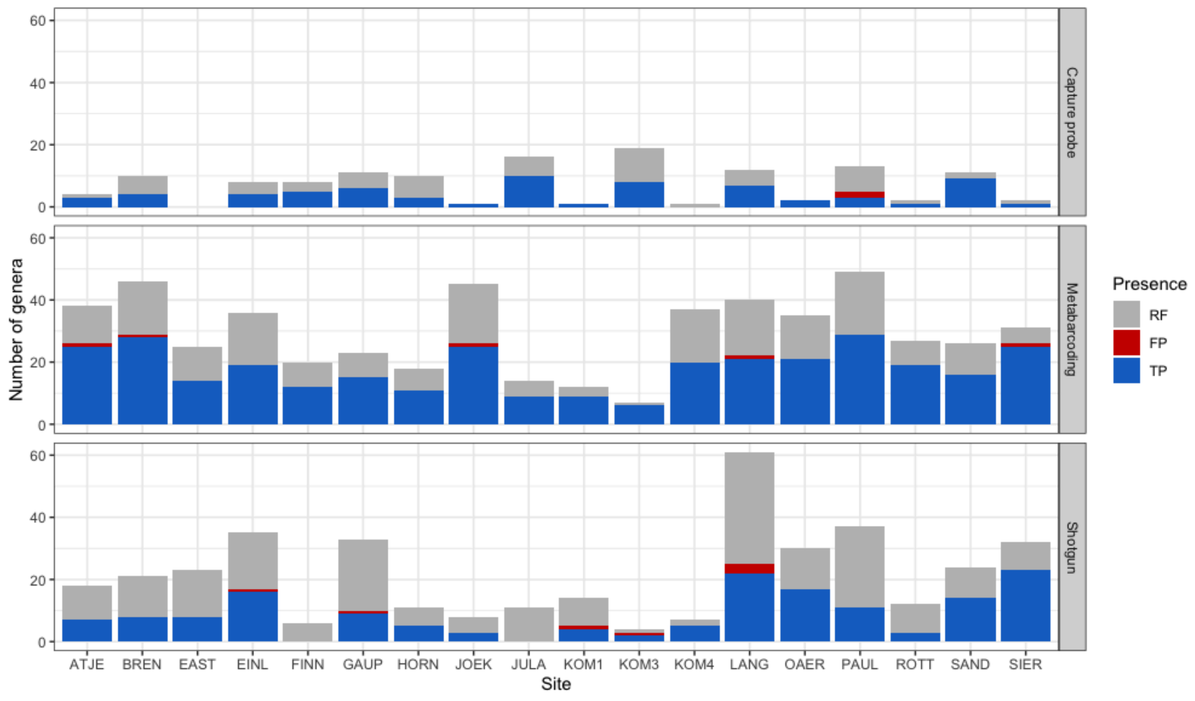


Figure S4. Taxonomic richness in different lakes detected by different methods based on optimal filtering.

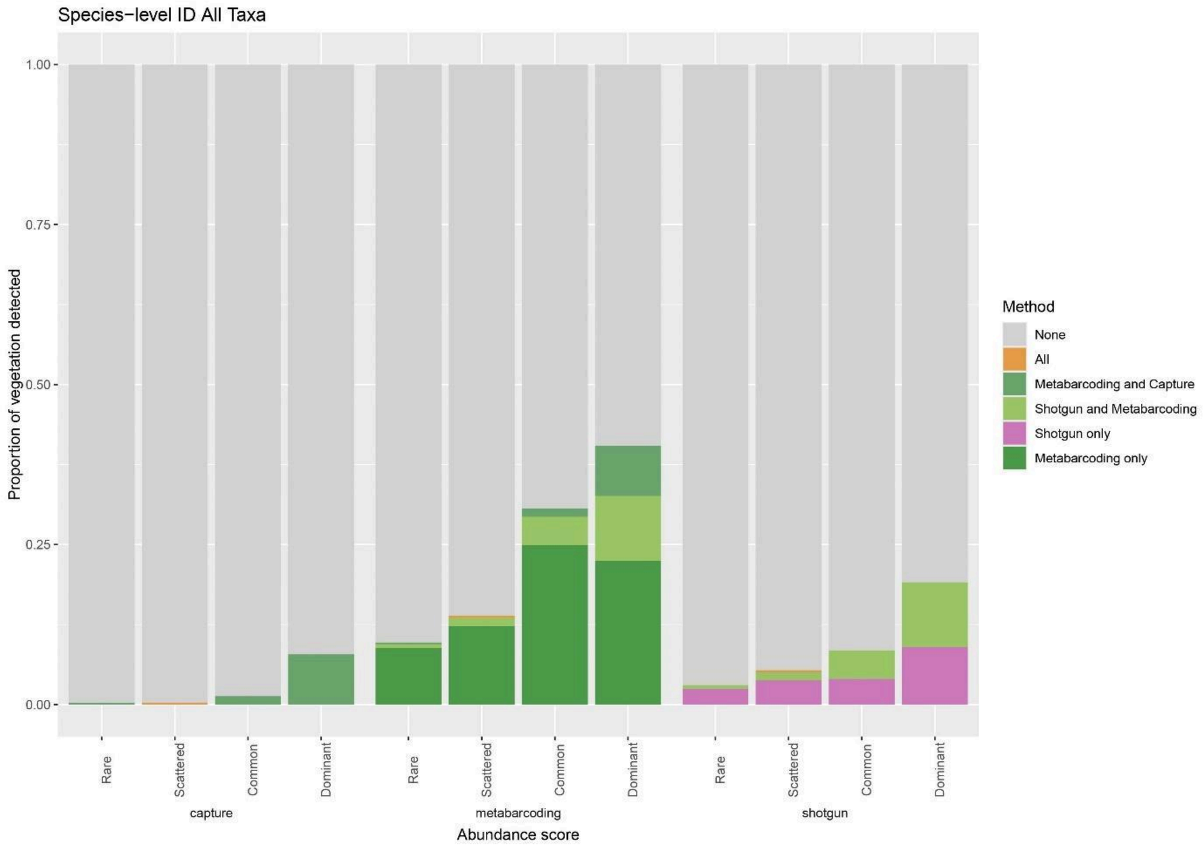


Figure S5. The proportion of 2 m vegetation survey taxa with a DNA match at the species-level grouped by DNA method detected by and ranked abundance. The DNA data shown are those done after the final optimal filtering.

### Supplementary Tables

For supplementary tables S1 - S5, see comparison_supplement.xlsx.

Table S6. Results from ordinal logistic regressions using the proportion of reads to predict plant abundance categories at the species and at the genus level *(matches, e.g., species and genus level matches between each DNA method and the vegetation surveys).* The data are filtered based on the optimal approach. The ranked abundances are only for the <2m vegetation surveys. In addition to raw read proportions, three different transformations of the data were evaluated: log, square root and double root. For each model, coefficients are shown with standard error (Est ± SE), p-values and ∆AIC, the difference in AIC to the model with the lowest AIC value. Models with the lowest AIC values and a significant relationship between read proportion and abundance category are highlighted in bold. Models with ∆AIC < 2 that are considered equally suitable are highlighted in italics. “f” indicates that models with a flexible threshold were preferred over models with an equidistant threshold “e” between the categories.

| **Method** | **Transf.** | **Species level** | | | **Genus level** | | |
|---|---|---|---|---|---|---|---|
| | | **Est ± SE** | **p** | **∆AIC** | **Est ± SE** | **p** | **∆AIC** |
| Capture probe<br><br>n spec. = 12<br><br>n gen. = 56 | none | -1.553 ± 1.139, e | 0.173 | 0 | -0.035 ± 0.230, e | 0.879 | 1.58 |
| | log | -2.564 ± 2.151, e | 0.233 | 0.44 | 0.067 ± 0.267, e | 0.802 | 0.06 |
| | sqrt | -1.514 ± 1.101, e | 0.169 | 0.20 | -0.003 ± 0.241, e | 0.989 | 0 |
| | double sqrt | -1.709 ± 1.312, e | 0.193 | 0.30 | 0.032 ± 0.251, e | 0.900 | 0.405 |
| Metabarcoding<br><br>n spec. = 178<br><br>n gen. = 346 | none | 0.615 ± 0.204, f | 0.003 | 21.85 | 0.677 ± 0.159, e | <0.001 | 27.01 |
| | *log* | *0.805 ± 0.148, f* | *<0.001* | *1.43* | 0.749 ± 0.110, e | <0.001 | 4.77 |
| | sqrt | 0.832 ± 0.193, f | <0.001 | 6.44 | 0.798 ± 0.134, e | <0.001 | 8.21 |
| | **double sqrt** | **0.859 ± 0.163, f** | **<0.001** | **0** | **0.817 ± 0.119, e** | **<0.001** | **0** |
| Shotgun<br><br>n spec. = 75<br><br>n gen. = 163 | none | 1.187 ± 0.524, e | 0.023 | 1.16 | 0.498 ± 0.184, e | 0.007 | 4.17 |
| | log | 0.723± 0.228, e | 0.001 | 0.62 | *0.554 ± 0.169, e* | *0.001* | *0.95* |
| | sqrt | 0.820 ± 0.280, e | 0.003 | 0.06 | *0.558 ± 0.172, e* | *0.001* | *0.24* |
| | double sqrt | 0.767 ±0.239, e | 0.002 | 0 | **0.568 ± 0.169, e** | **0.001** | **0** |

Table S7. Results from ordinal logistic regressions using the read numbers to predict plant abundance categories at the species and at the genus level *(matches, e.g., species and genus level matches between each DNA method and the vegetation surveys).* The data are filtered based on the optimal approach. The ranked abundances are only for the <2m vegetation surveys. In addition to raw read numbers, three different transformations of the data were evaluated: log, square root and double root. For each model, coefficients are shown with standard error (Est ± SE), Sp-values and ∆AIC, the difference in AIC to the model with the lowest AIC value. Models with the lowest AIC values and a significant relationship between read proportion and abundance category are highlighted in bold. Models with ∆AIC < 2 that are considered equally suitable are highlighted in italics. "f" indicates that models with a flexible threshold were preferred over models with an equidistant threshold "e" between the categories.

| **Method** | **Transf.** | **Species level** | | | **Genus level** | | |
|---|---|---|---|---|---|---|---|
| | | **Est ± SE** | **p** | **∆AIC** | **Est ± SE** | **p** | **∆AIC** |
| Capture probe<br>n spec. = 12<br>n gen. = 56 | none | -1.239 ± 1.232, e | 0.315 | 0 | -0.221 ± 0.208, e | 0.287 | 1.10 |
| | log | 0.154 ± 0.872, e | 0.860 | 1.93 | -0.002 ± 0.247, e | 0.992 | 0.57 |
| | sqrt | -0.839 ± 1.088, e | 0.441 | 1.08 | -0.174 ± 0.224, e | 0.438 | 0 |
| | double sqrt | -0.413 ± 0.996, e | 0.669 | 1.75 | -0.101 ± 0.237, e | 0.669 | 0.40 |
| Metabarcoding<br>n spec. = 197<br>n gen. = 346 | none | 0.647 ± 0.222, f | <0.001 | 19.29 | 0.766 ± 0.175, e | <0.001 | 19.53 |
| | *log* | *0.777 ± 0.154, f* | *<0.001* | *1.14* | 0.713 ± 0.110, e | <0.001 | 7.75 |
| | sqrt | 0.767 ± 0.181, f | <0.001 | 4.75 | 0.823 ± 0.137, e | <0.001 | 4.10 |
| | **double sqrt** | **0.814 ± 0.163, f** | **<0.001** | **0** | **0.798 ± 0.118, e** | **<0.001** | **0** |
| Shotgun<br>n spec. = 93<br>n gen. = 163 | none | 2.143 ± 1.845, e | 0.245 | 2.36 | 0.273 ± 0.154, e | 0.076 | 6.52 |
| | **log** | **0.553± 0.216, e** | **0.010** | **0** | **0.461 ± 0.161, e** | **0.004** | **0** |
| | sqrt | 0.788 ± 0.185, e | 0.075 | 0.84 | 0.354 ± 0.160, e | 0.027 | 3.26 |
| | *double sqrt* | *0.602 ±0.261, e* | *0.021* | *0.19* | *0.414 ± 0.160, e* | *0.010* | *1.48* |
